# Inhalable Nanobody (PiN-21) prevents and treats SARS-CoV-2 infections in Syrian hamsters at ultra-low doses

**DOI:** 10.1101/2021.02.23.432569

**Authors:** Sham Nambulli, Yufei Xiang, Natasha L. Tilston-Lunel, Linda J. Rennick, Zhe Sang, William B. Klimstra, Douglas S. Reed, Nicholas A. Crossland, Yi Shi, W. Paul Duprex

**Author notes:** Corresponding author (Y.S.) or (W.P.D.). Equal contributions (Sham Nambulli, Yufei Xiang).

## Abstract

Globally there is an urgency to develop effective, low-cost therapeutic interventions for coronavirus disease 2019 (COVID-19). We previously generated the stable and ultrapotent homotrimeric Pittsburgh inhalable Nanobody 21 (PiN-21). Using Syrian hamsters that model moderate to severe COVID-19 disease, we demonstrate the high efficacy of PiN-21 to prevent and treat SARS-CoV-2 infection. Intranasal delivery of PiN-21 at 0.6 mg/kg protects infected animals from weight loss and substantially reduces viral burdens in both lower and upper airways compared to control. Aerosol delivery of PiN-21 facilitates deposition throughout the respiratory tract and dose minimization to 0.2 mg/kg. Inhalation treatment quickly reverses animals’ weight loss post-infection and decreases lung viral titers by 6 logs leading to drastically mitigated lung pathology and prevents viral pneumonia. Combined with the marked stability and low production cost, this novel therapy may provide a convenient and cost-effective option to mitigate the ongoing pandemic.

## Introduction

By January 2021, a year after the outbreak of severe acute respiratory syndrome coronavirus 2 (SARS-CoV-2) was first reported (*1*), close to 100 million people have been infected by this highly transmissible virus resulting in significant morbidity and mortality worldwide. In addition to vaccines, there is an unparalleled quest to develop innovative and cost-effective therapeutics to combat the COVID-19 pandemic (*2, 3*). Early treatment using high-titer convalescent plasma (CP) may reduce the risk of severe disease in seniors (*4*), although CP is limited by supply. Potent neutralizing monoclonal antibodies (mAbs), predominantly isolated from COVID-19 patients for recombinant productions, have been developed for passive immunotherapy (*5–14*). *In vivo* evaluations of mAbs in animal models of COVID-19 disease such as murine, hamster, and nonhuman primates (NHPs), have provided critical insights into efficacy and the mechanisms by which they alter the course of infection (*15–24*). While mAb therapy lifts hopes to treat mild symptom onset in patients, they nevertheless require exceedingly high administration doses-typically several grams for intravenous (*i.v.*) injection (*25, 26*). The requirement of high doses for efficient neutralization may reflect SARS-CoV-2 virulence, pathogenesis, and the notoriously low efficiency of *i.v.* delivering these relatively large biomolecules across the plasma-lung barrier to treat pulmonary infections (*27*). Moreover, the associated high costs and challenges in bulk manufacturing may further limit the broad clinical use of mAbs worldwide (*2*).

In parallel efforts, we and others have recently developed camelid single-domain antibody fragments or nanobodies (Nbs) that primarily target the receptor-binding domain (RBD) of the SARS-CoV-2 spike (S) glycoprotein for virus neutralization (*14, 28–31*). Highly selected Nbs and the multivalent forms obtain high neutralization potency comparable to, or even better (per-mass) than, some of the most successful SARS-CoV-2 neutralizing mAbs. In particular, an ultrapotent homotrimeric construct, Pittsburgh inhalable Nanobody 21 (PiN-21), efficiently blocked SARS-CoV-2 infectivity at below 0.1 ng/ml *in vitro* (*28*). Compared to mAbs, Nbs are substantially cheaper to produce. Moreover, affinity-matured, ultrapotent Nbs are characterized by high solubility and stability that facilitate drug scaling, storage, and transportation, all of which are critical in response to pandemics. The excellent physicochemical properties and small sizes of Nbs raise an exciting possibility of efficient pulmonary delivery by aerosolization with characteristics of rapid onset of action, high local drug concentration/bioavailability, and improved patient compliance (needle-free) that may benefit a large population of SARS-CoV-2 infected patients (*27–29, 32*). However, despite the promise, no successful *in vivo* studies have been reported to date. The inferior pharmacokinetics of monomeric Nbs due to their small size (~15 kDa) and a lack of Fc-mediated immune effectors’ function, which is often required to augment the *in vivo* neutralizing activities of mAbs (*33–35*), drive potential concerns for Nb-based therapy. It remains unknown if the high *in vitro* neutralization potency of SARS-CoV-2 Nbs can be translated into *in vivo* therapeutic benefits.

In this study, we systematically evaluated the efficacy of PiN-21 for prophylaxis and treatment of SARS-CoV-2 infected Syrian hamsters which model moderate to severe COVID-19 disease. We provided direct evidence that ultra-low administration of PiN-21 efficiently treats the virus infection. Notably, PiN-21 aerosols can be inhaled to target respiratory infection which drastically reduces viral loads and prevents lung damage and viral pneumonia. This novel Nb-based therapy shows high potentials for the treatment of early infection and may provide a robust and affordable solution to address the current health crisis.

## Results

### PiN21 efficiently protects and treats SARS-CoV-2 infection in Syrian hamsters

To assess the *in vivo* efficacy of PiN-21, 12 hamsters were divided into two groups and infected with 9 x 10^4^ plaque-forming units (p.f.u.) of SARS-CoV-2 *via* the intratracheal (IT) route. Shortly after infection, Nb was delivered intranasally (IN) at an average dose of 0.6 mg/kg (**Fig 1A**). Animals were monitored daily for weight change and clinical signs of disease. Half of the animals were euthanized 5 days post-infection (d.p.i.) and the remaining were euthanized 10 d.p.i. Virus titers in lung samples from the euthanized animals were measured by plaque assay. Nasal washes and throat swabs were collected at 2 and 4 d.p.i. to determine viral loads in the upper respiratory tract. Consistent with published studies (*36, 37*), IT inoculation of hamsters with SARS-CoV-2 resulted in a robust infection, rapid weight loss in all animals up to 16% at 7 d.p.i. and resulting recovery and reversal of weight loss by 10 d.p.i. before recovery. However, concurrent IN delivery of PiN-21 eliminated any significant weight loss in the infected animals (**Fig 1B**). This dramatic protection was accompanied by a reduction of viral titer in the lungs, with an average decrease of 4 order of magnitude in the lung tissue, respectively, compared to control on 5 d.p.i. (**Fig 1C**). Consistently, a 3-log reduction of the viral genomic RNA (gRNA) by reverse transcriptase (RT)-qPCR was evident on 5 and 10 d.p.i. (**Fig S1A-B**). Infectious virus was essentially cleared by 10 d.p.i.

**Figure 1.**
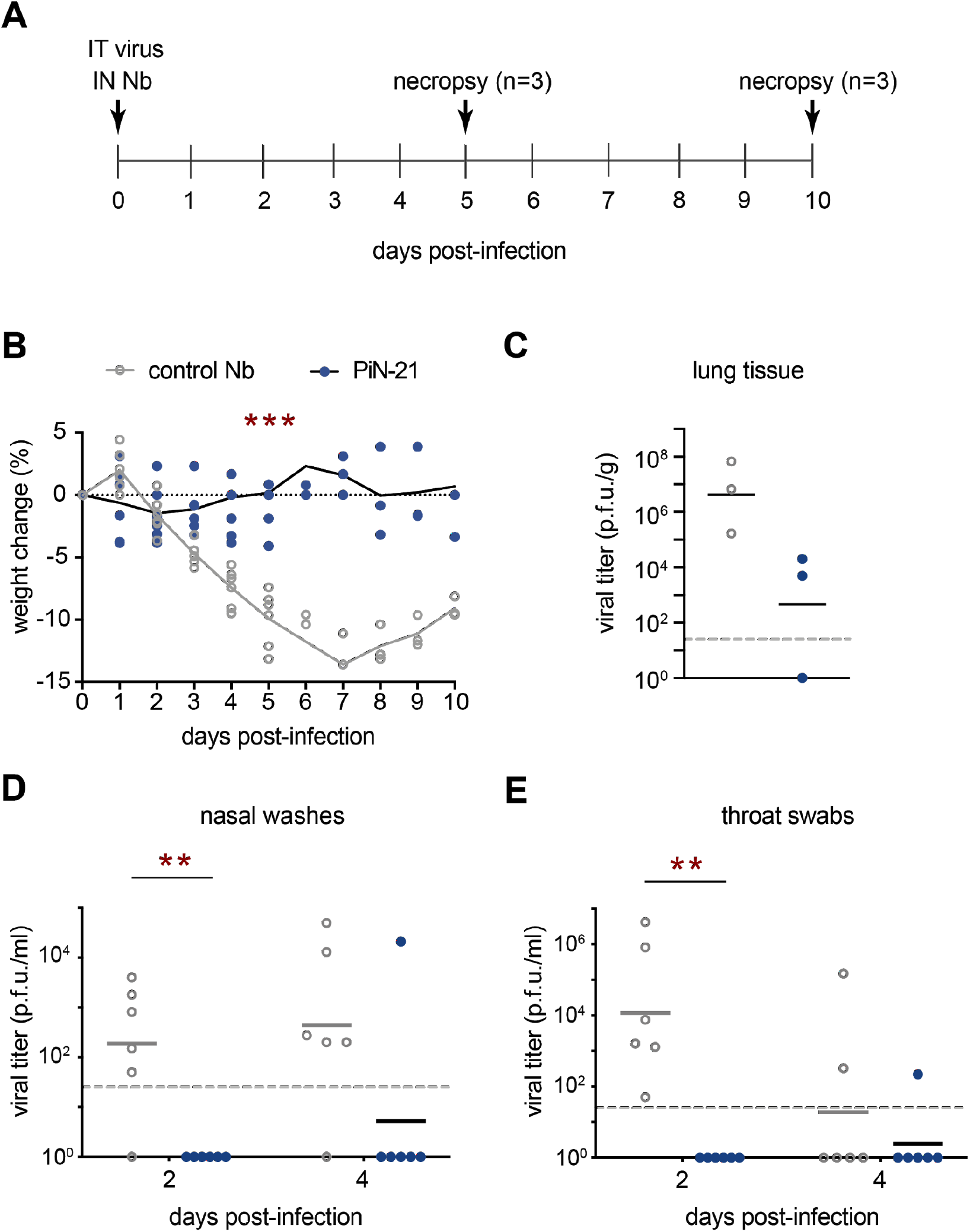
PiN-21 protects Syrian hamsters from SARS-CoV-2 infection. **A.** Overview of the experimental design. 9 x 10^4^ p.f.u. of SARS-CoV-2 was IT inoculated followed by IN delivery of 100 μg PiN-21 or a control Nb. Animal weight changes were monitored daily. Nasal washes and throat swabs were collected on 2 and 4 d.p.i. Animals were euthanized for necropsy on 5 (n=3) and 10 d.p.i (n=3). **B.** The protection of weight loss of infected hamsters treated with PiN-21. *** indicates a p-value of < 0.001. **C-E.** Measurement of viral titers by the plaque assay. ** indicates a p-value of < 0.01. The dashed line indicates the detection limit of the assay.

Notably, the virus was undetectable in the upper respiratory tract (URT) including both nasal washes and throat swabs of all PiN-21-treated animals on 2 d.p.i. This is significantly different from the control group, where varying levels of infectious virus were present (**Fig 1D-E**). Furthermore, five out of six PiN-21 treated animals remained protected from detectable infection 4 d.p.i. The results were further supported by a drastic decrease of gRNA in the URT (**Fig S1C-D**). Together, this demonstrates that the high *in vitro* neutralization potency of PiN-21 can be translated into therapeutic benefits *in vivo* independent of Fc-mediated immune responses. PiN-21 can efficiently protect SARS-CoV-2 infection in hamsters by rapidly and drastically suppressing viral replication in both the URT and lower respiratory tract (LRT).

Previous studies reveal that clinical mAbs are less effective for COVID-19 treatment (post-infection) than for prophylaxis (pre-infection) in animal models, possibly reflecting the virulence of SARS-CoV-2, speed of virus replication, and rapid symptom onset (*15, 22, 38*). Therefore, we evaluated the therapeutic potential of PiN-21 since it was highly effective when co-administered. To explore the second route of infection, hamsters were inoculated IN with 3 x 10^4^ p.f.u. of SARS-CoV-2. PiN-21 or a control Nb (0.6 mg/kg) was IN-delivered to animals 6 hours post-infection (h.p.i.). Animal weights were monitored daily, throat swabs and nasal washes were collected, before euthanized on 6 d.p.i. (**Fig S2A**). Similar to the IT route, IN-infection of hamsters with SARS-CoV-2 resulted in precipitous weight losses in the control animals. Encouragingly, intranasal treatment using PiN-21 significantly reduced weight loss throughout the assessment period (**Fig S2B**), paralleling the results of clinical mAbs in the same model albeit using substantially higher doses. Less than 100-fold reduction in virus titers were found in nasal washes and throat swabs on 2 and 4 d.p.i. (**Fig S2C-D**). Moreover, infectivity was undetectable in lung tissues 6 d.p.i., indicating that the virus has been predominantly cleared. Analysis of early time points will be needed to better understand virus suppression by Nb treatment.

### PiN21 aerosolization effectively treats SARS-CoV-2 infected hamsters at an ultra-low dose

The marked physicochemical properties of PiN-21 prompted us to evaluate pulmonary delivery by inhalation. To evaluate the impact of construct size and pharmacokinetics on lung uptake, we fused monomeric Nb21 and PiN-21 to an Nb that binds serum albumin (Alb) of both human and rodents with high affinity to generate two serum-stable constructs (Nb-21_Alb_ and PiN-21_Alb_) (*39*). Using a portable mesh nebulizer, we aerosolized Nb21_Alb_, PiN-21, and PiN-21_Alb_ and evaluated their post-aerosolization neutralization activities by pseudovirus neutralization assay. All constructs retained high neutralization potency *in vitro* (**Fig S3B**). The amount of Nbs recovered post-aerosolization was inversely correlated with the size of constructs (**Fig S3A**). Moreover, while Nb-21_Alb_ had the highest recovery, the post-aerosolization *in vitro* neutralization activity was substantially lower than for other constructs, Nb-21_Alb_ was therefore excluded from downstream therapeutic analyses.

Next, we compared two ultrapotent constructs PiN-21 and PiN-21_Alb_ for targeted aerosolization delivery into hamsters. Nbs were aerosolized using a nebulizer (Aerogen, Solo) that produces small aerosol particles with a mass median aerodynamic diameter of ~ 3 μm (**Table S1**). Animals were sacrificed 8 and 24 hours post-administration to assess Nb distribution and activities when recovered from various respiratory compartments and sera (**Fig 2A**). Consistent with the result using the portable nebulizer, we found the inhalation dose of PiN-21 was approximately two times PiN-21_Alb_ (at 8 h, 41.0 μg or 0.24 mg/kg for PiN-21 v.s. 23.7 μg or 0.13 mg/kg for PiN-21_Alb_) (**Table S1**) while neutralization activity as assessed by plaque reduction neutralization test (PRNT_50_) of SARS-CoV-2 remained essentially unchanged after aerosolization of both Nb constructs (**Fig 2B**).

**Figure 2.**
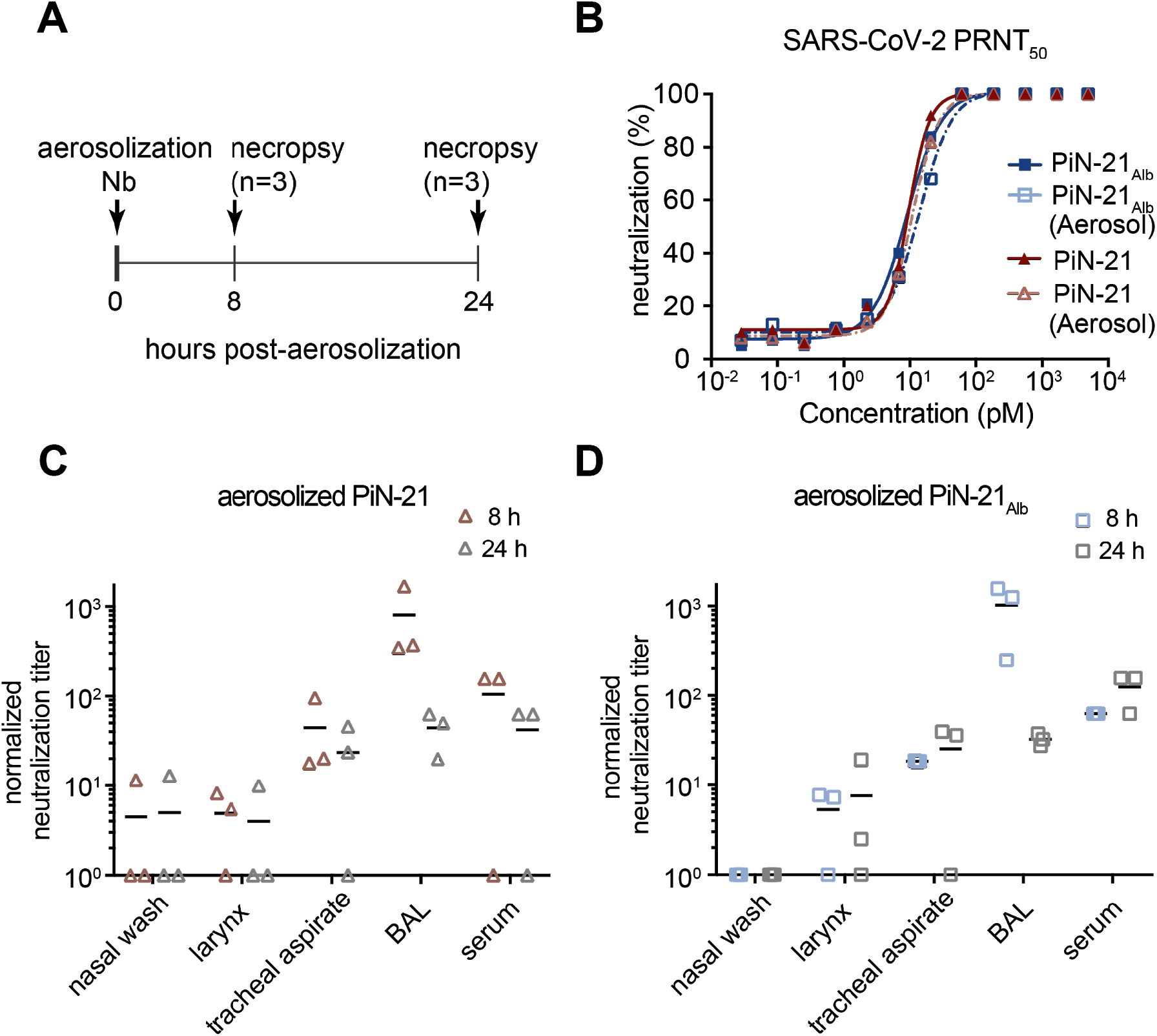
Assessment of Nb delivery in the hamster respiratory system. **A.** Schematic design of PiN-21 and PiN-21_Alb_ aerosolization in hamster models. **B.** Nb neutralization potency before and after aerosolization measured by PRNT50 assay. **C and D.** Normalized overall neutralization activity by plaque assay of PiN-21 and PiN-21_Alb_ of different time points post-aerosolization.

The neutralization activities of both Nb constructs were detected throughout the respiratory tract and in sera. As expected, within the airways the neutralizing activities were predominantly associated with bronchoalveolar lavage (BAL) fluid, followed by tracheal aspirate, larynx wash, and nasal wash samples (**Fig 2C-D**). Compared to 8-hour post-inhalation, we found that the amounts and activities of Nbs in BAL, but not in sera, were substantially lower 24-hour post-inhalation, possibly indicating more rapid clearance. In addition, Nb conjugation to serum albumin did not seem to impact the activities in the airways, whilst stability was enhanced in the serum. These data underscore the requirement of an ultra-low dose of the ultrapotent PiN-21 construct to neutralize SARS-CoV-2 infectivity *in vivo* efficiently. Finally, PiN-21 was preferentially selected for further evaluation owing to the high stability and resistance to aerosolization, which are likely critical for clinical applications.

To assess the therapeutic efficacy of PiN-21 by inhalation, 12 hamsters were IN inoculated with SARS-CoV-2 (3 x 10^4^ p.f.u.) followed by single-dose aerosolization treatment of either PiN-21 or the control Nb at 6 h.p.i. Animals were monitored for weight loss, throat swabs and nasal washes were collected daily. Animals were euthanized (3 d.p.i.) and lungs and trachea were collected for virological, histopathology, and immunohistochemical analysis (**Fig 3A**). Notably, pulmonary delivery of PiN-21 aerosols, despite only a *minute* amount, led to a remarkable reverse of weight loss in the treated animals. The average weight gain was 2% in PiN-21 versus 5% loss in the control on 3 d.p.i. (**Fig 3B**). The weight loss in the control group was highly reproducible when compared with the above experiments. Critically, aerosolization treatment diminished infectious viruses in lung tissue by 6 orders of magnitude (**Fig 3C**). The treatment also substantially decreased virus gRNA in the lungs (**Fig S4C**). Moreover, we observed a substantial reduction of viral titers in nasal washes and throat swabs (**Fig S4A-B**). This indicates that Nb administration by aerosolization may limit human-to-human transmission of SARS-CoV-2.

**Figure 3.**
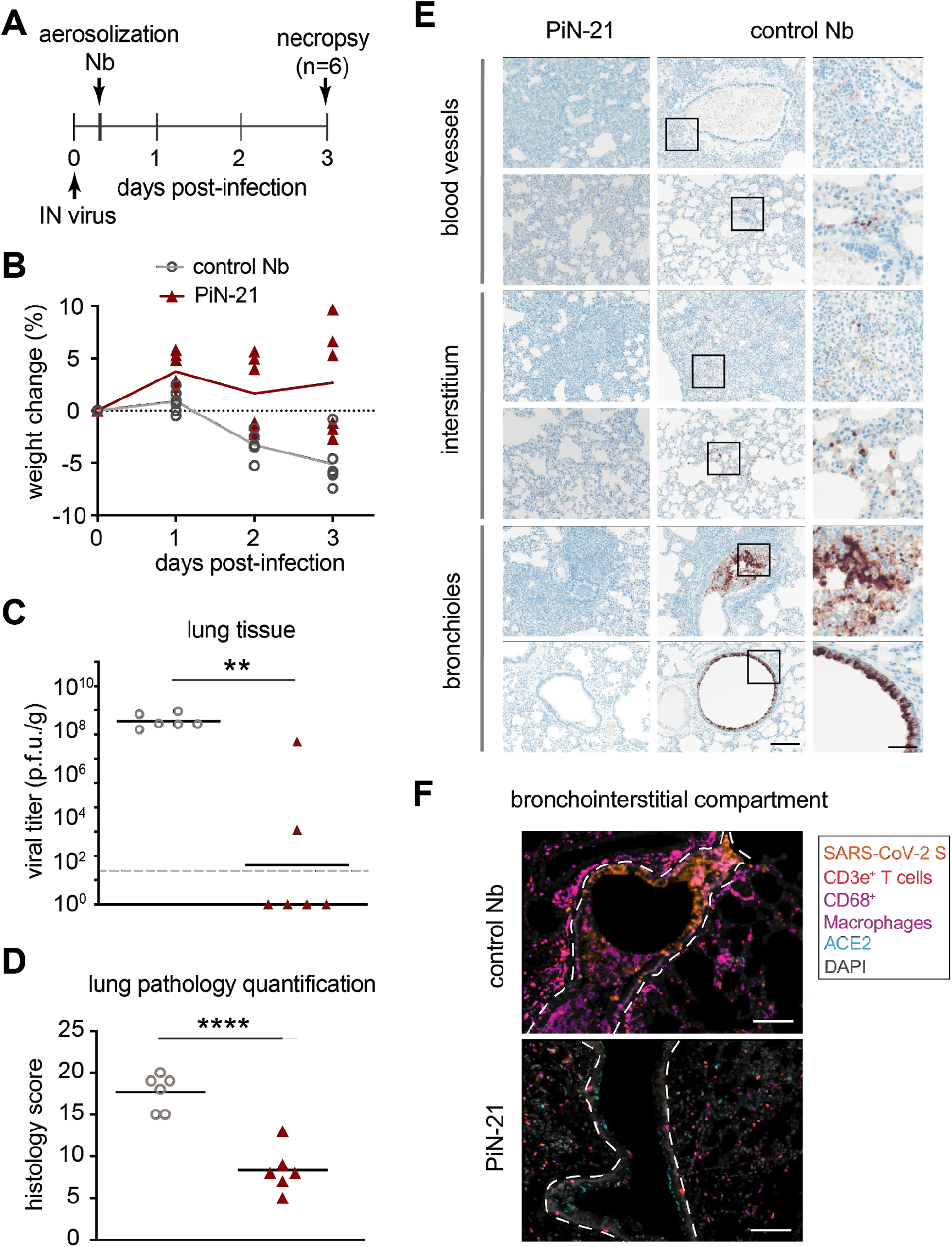
Treatment efficacy of aerosolized PiN-21 in the hamster model of SARS-CoV-2. **A.** Overview of the experiment design. 3 x 10^4^ p.f.u. of SARS-CoV-2 was IN inoculated. PiN-21 or a control Nb was aerosolized to hamsters in the cage 6 h.p.i. Animal weight changes were monitored, nasal washes and throat swabs were taken daily. Animals were euthanized for necropsy on 3 d.p.i. **B.** The percentage of body weight change of PiN-21 aerosol-treated animals compared to the control (n=6). **C.** Reduction of viral titers in hamster lungs (3 d.p.i.). Significant differences were observed between treated and control groups. **, P < 0.01; *, P < 0.05. The dashed line indicates the detection limit of the assay. **D.** Lung pathology scores of treated and control groups. Significant difference was denoted by ****, P < 0.0001. **E.** H&E staining of necrotizing bronchointerstitial pneumonia affiliate with abundant SARS-CoV-2 S antigen in bronchiole epithelium and alveolar type 1 and 2 pneumocytes in the control group. All images acquired at 20 x, scale bar = 100 μm. Areas marked by boxes are shown at higher magnification in the right-most panel (scale bar = 25 μm). **F.** Immunostainings of bronchointerstitial compartments (3 d.p.i.). Orange, SARS-CoV-2 S; magenta, CD68/macrophages; red, CD3e+ T cells; teal, ACE2; Grey, DAPI. The bronchiole is outlined by white hash. Total magnification 200x, scale bar = 100 μm.

### Effective control of SARS-CoV-2 infection in the LRT of PiN-21 aerosolized animals

To understand the mechanisms by which Nb aerosols prevent and/or ameliorate lower respiratory disease caused by SARS-CoV-2 infection better, we performed whole lung semi-quantitative ordinal histologic analysis of control (n=6) and PiN-21 (n=6) treated animals euthanized at 3 d.p.i (**Table S2-3**). Cumulative scores encompassed pathologic features of airways, blood vessels, and alveoli/pulmonary interstitium. PiN-21 aerosolization protected most animals (5/6) from severe COVID-related histopathologic disease reflected by decreased ordinal scores (P < 0.0001) when compared to Nb treated controls (**Fig 3D, Table S4**). Histopathologic findings in the control group resembled previous reports of SARS-CoV-2 inoculation in Syrian hamsters (*40, 41*). Pulmonary disease observed in the PiN-21 treated animals was very mild (**Fig 3E, Fig S5**) being characterized by the absence of severe necrotizing bronchiolitis in the majority of animals (5/6), a pathological finding ubiquitously observed in all control animals. Furthermore, the single PiN-21 treated animal with necrotizing bronchiolitis had localized disease, compared to the multifocal and bilateral distribution observed in most Nb controls (4/6). Bronchiolitis was also affiliated with less severe bronchial hyperplasia and hypertrophy and absence of syncytial cells when compared to Nb controls. The predominant histologic finding in PiN-21 treated animals was minimal-to-mild perivascular and peribronchial mononuclear inflammation consisting of macrophages and lymphocytes. Furthermore, aside from the one animal already mentioned, the PiN-21 group had considerably less interstitial inflammation with decreased vascular permeability, as indicated by the absence of perivascular and intra-alveolar edema, hemorrhage, and fibrin exudation (**Fig S5**).

In control animals, S antigen was abundant in the cytoplasm of the bronchiolar epithelium, with less common detection in alveolar type 1 and 2 pneumocytes. Interstitial and peribronchiolar infiltrates were composed of large numbers of CD3e+ T cells and CD68+ macrophages, with a complete absence of angiotensin-converting enzyme 2 (ACE2) in the apical cytoplasm of bronchiole epithelium in areas with abundant viral S (**Fig 3F, upper panel**). Consistent with a striking 6-log virus reduction after aerosolization, S antigen was extremely sparse (<1% of permissive cells) in all PiN-21 treated animals, with decreased T cell and macrophage immune cell infiltrate, and retention of native apical bronchiole ACE2 expression (**Fig 3F, lower panel**).

To determine the impact of PiN-21 on the upper airways of the lower respiratory tract we also examined the trachea histologically. In PiN-21 treated animals, tracheas for all animals were within normal limits, while mild to moderate neutrophilic and lymphohistiocytic tracheitis, with variable degrees of degeneration and necrosis, and segmental hyperplasia and hypertrophy were observed in all the control animals (**Fig 3F**). In summary, our data clearly demonstrate that PiN-21 aerosolization given during early disease course is highly effective in decreasing SARS-CoV-2 entry, subsequent replication in permissive epithelial cells of the lower respiratory tract and this, in turn, has a major impact on viral shedding. The result is the prevention of disease, including decreased cytopathic effect on permissive epithelial cells, retention of ACE2 expression on permissive bronchioles, and decreased recruitment of inflammatory cells to sites of replication.

## Discussion

In this work, we demonstrate the high therapeutic efficacy of a trimeric Nb against SARS-CoV-2 infection in Syrian hamsters. Our investigations leverage both intranasal and aerosol delivery of PiN-21 and demonstrate Nb treatment effectively targets the deep and local pulmonary structures such as terminal alveoli, which are lined with alveolar cells rich in ACE2 receptor to block viral entry and replication efficiently. Moreover, infection-induced weight loss correlates with pulmonary virus titer (Pearson r = −0.7) (**Fig S6**) and this clinical sign may be used to indicate the onset of infections (*13*). Notably, the ability of PiN-21 to eradicate viral replication and lung pathology almost completely in both the URT and LRT in hamsters contrasts the effects recently shown by clinical antibodies, which remain particularly challenging to treat SARS-CoV-2 infection in the same model (*15*).

Significantly improved delivery upon aerosolization may be anticipated in NHPs and humans since airway anatomical structures differ considerably from small rodents in which a high degree of inertial impaction is seen using liquid droplets. Several inhalation therapeutics with excellent safety profiles are commercially available and many are under clinical trials (*27, 42*). A combination of extremely low deposit doses will minimize potential adverse effects. We envision that PiN-21 aerosolization treatment could provide both a convenient and cost-effective solution to alleviate disease onset and reduce virus transmission, especially for mild COVID-19 patients who constitute major populations of infections. It may also benefit high-risk groups, such as seniors, immunocompromised individuals, and infants, in both inpatient and outpatient settings. Finally, as prevalent circulating variants of SARS-CoV-2 have emerged to evade clinical antibodies and wane vaccine-elicited serologic responses (*43–47*), this proof-of-concept study will shed light on the use of stable, multi-epitope and multivalent Nb constructs, in combination with PiN-21, as a novel aerosol cocktail to effectively block virus mutational escape (*28*).

## Supplemental Figures

**Figure S1.**
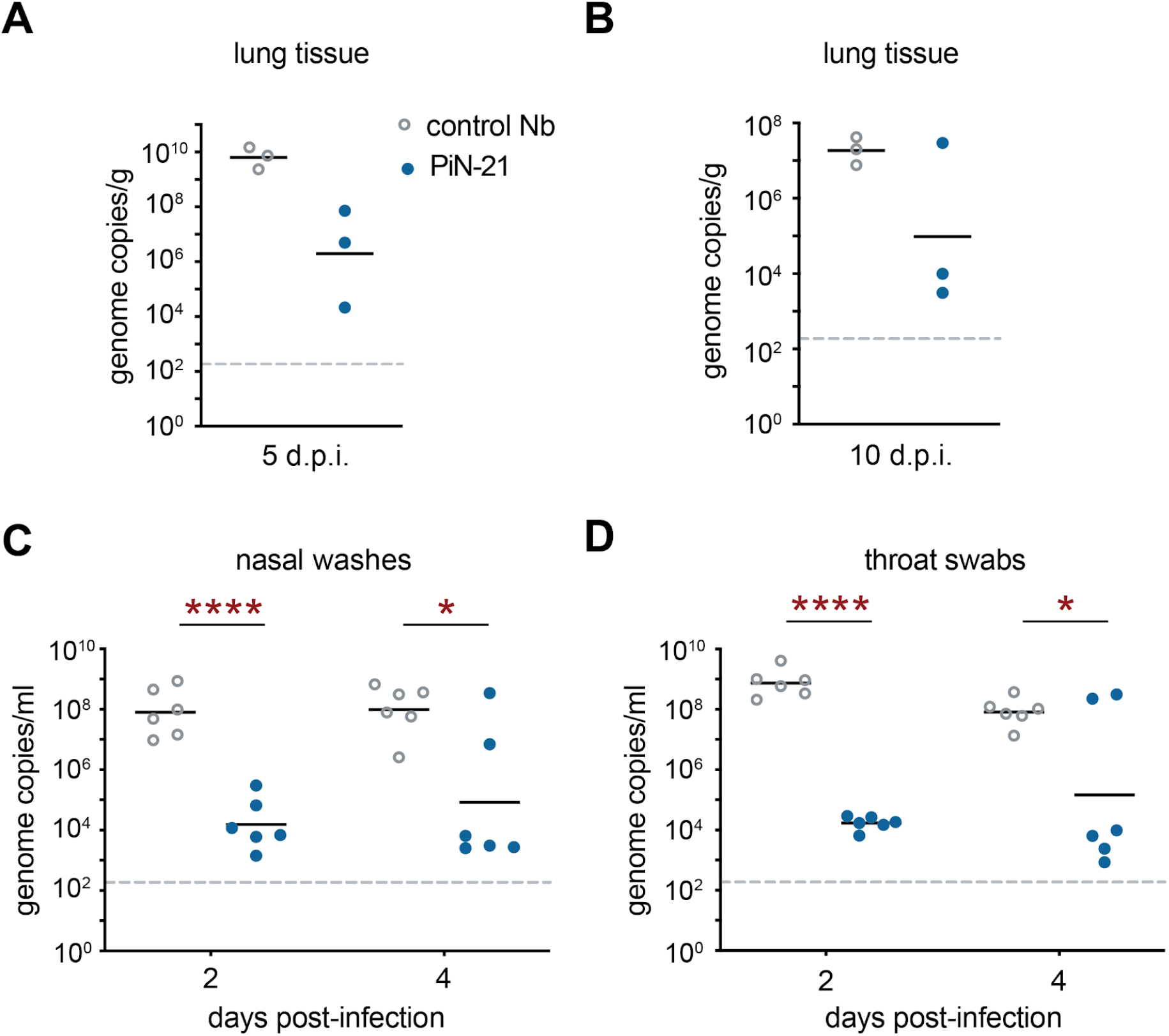
The efficacy of PiN-21 for protecting SARS-CoV-2 infection in hamsters. **A-B.** Measurement of gRNA by RT-qPCR on 5 and 10 d.p.i. (n = 3 in each group). **C-D.** Measurement of gRNA by RT-qPCR on 2 and 4 d.p.i. * indicates a p-value of < 0.05. **** indicates a p-value of < 0.0001. The dashed line indicates the detection limit of the assay.

**Figure S2.**
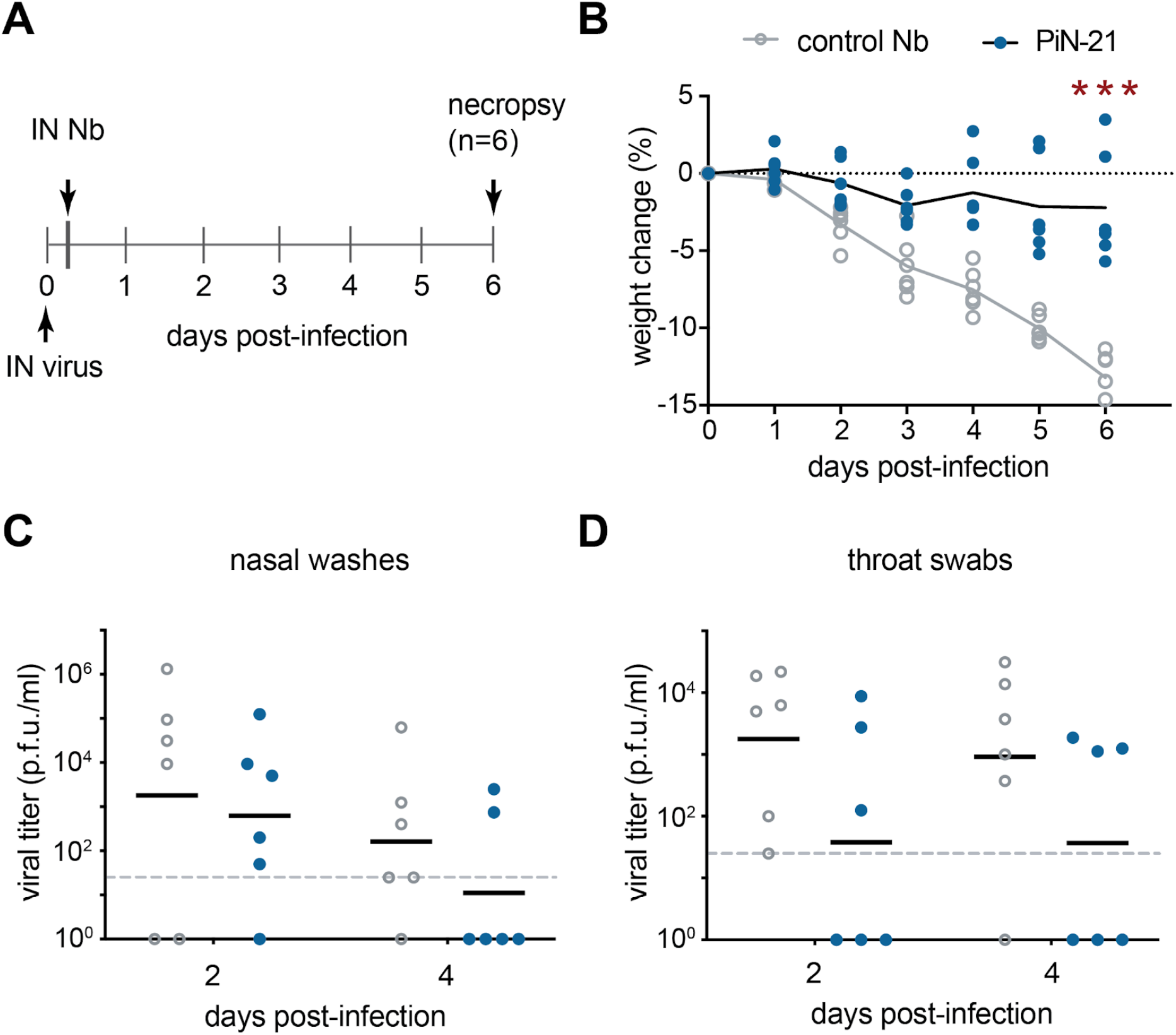
The efficacy of PiN-21 for the treatment of SARS-CoV-2 infection in hamsters. **A.** Schematic design of IN delivery of PiN-21 in hamsters for treatment. 3 x 10^4^ p.f.u. of SARS-CoV-2 was IN inoculated. 100 μg of PiN-21 or a control Nb was IN delivered 6 h.p.i. Animal weight changes were monitored daily and were euthanized for necropsy on 6 d.p.i. **B.** Protection of weight loss in the PiN-21 treatment group. Significant differences were denoted as **, P < 0.01; ***, P < 0.001. **A-B.** Measurement of viral titers in nasal washes and throat swabs by the plaque assay on 2 and 4 d.p.i. (n = 6).

**Figure S3.**
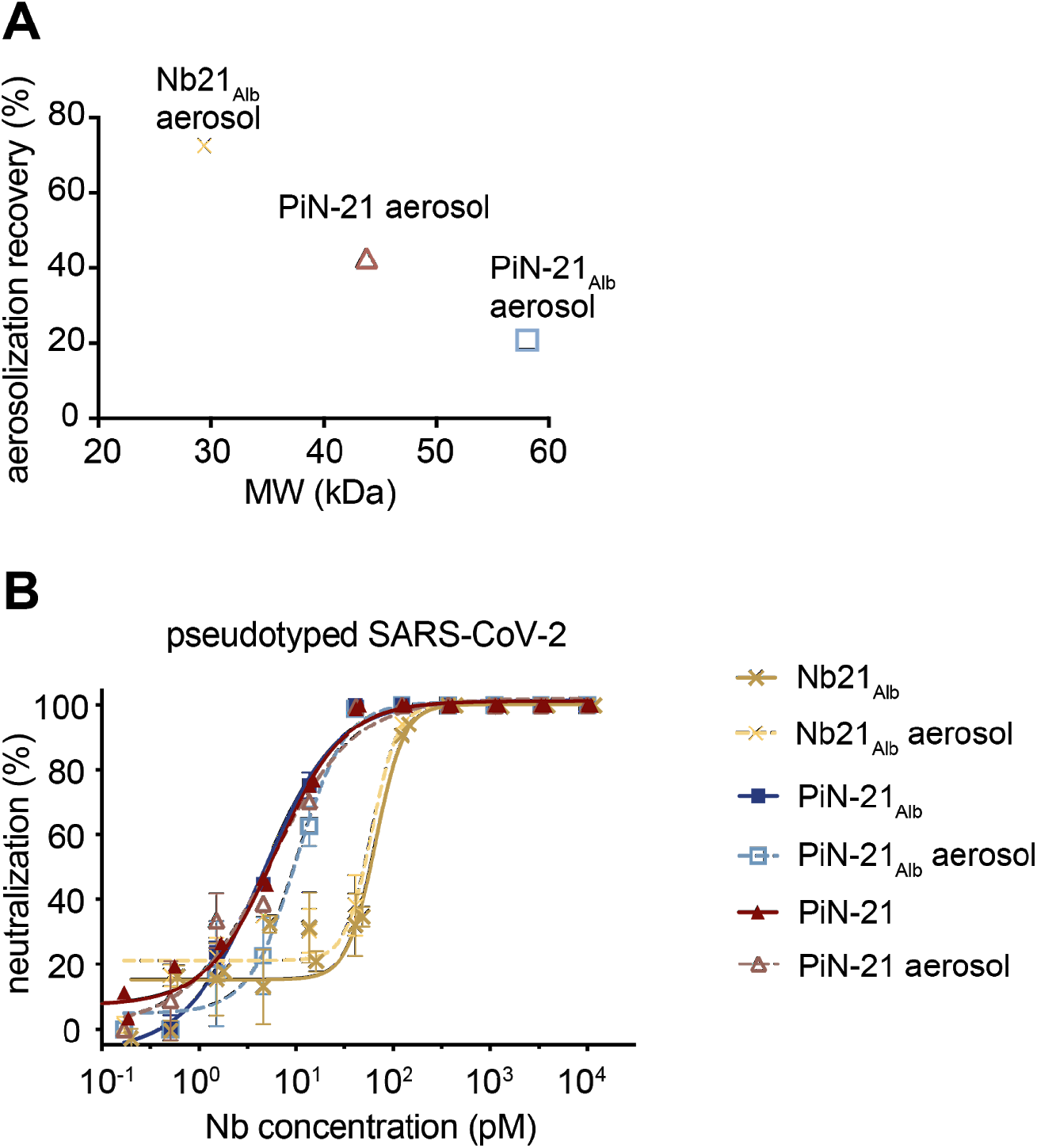
Characterization of Nb constructs after aerosolization by a portable mesh nebulizer. **A.** Protein recovery after aerosolization. **B.** Nb neutralization potency before and after aerosolization measured by pseudovirus neutralization assay.

**Figure S4.**
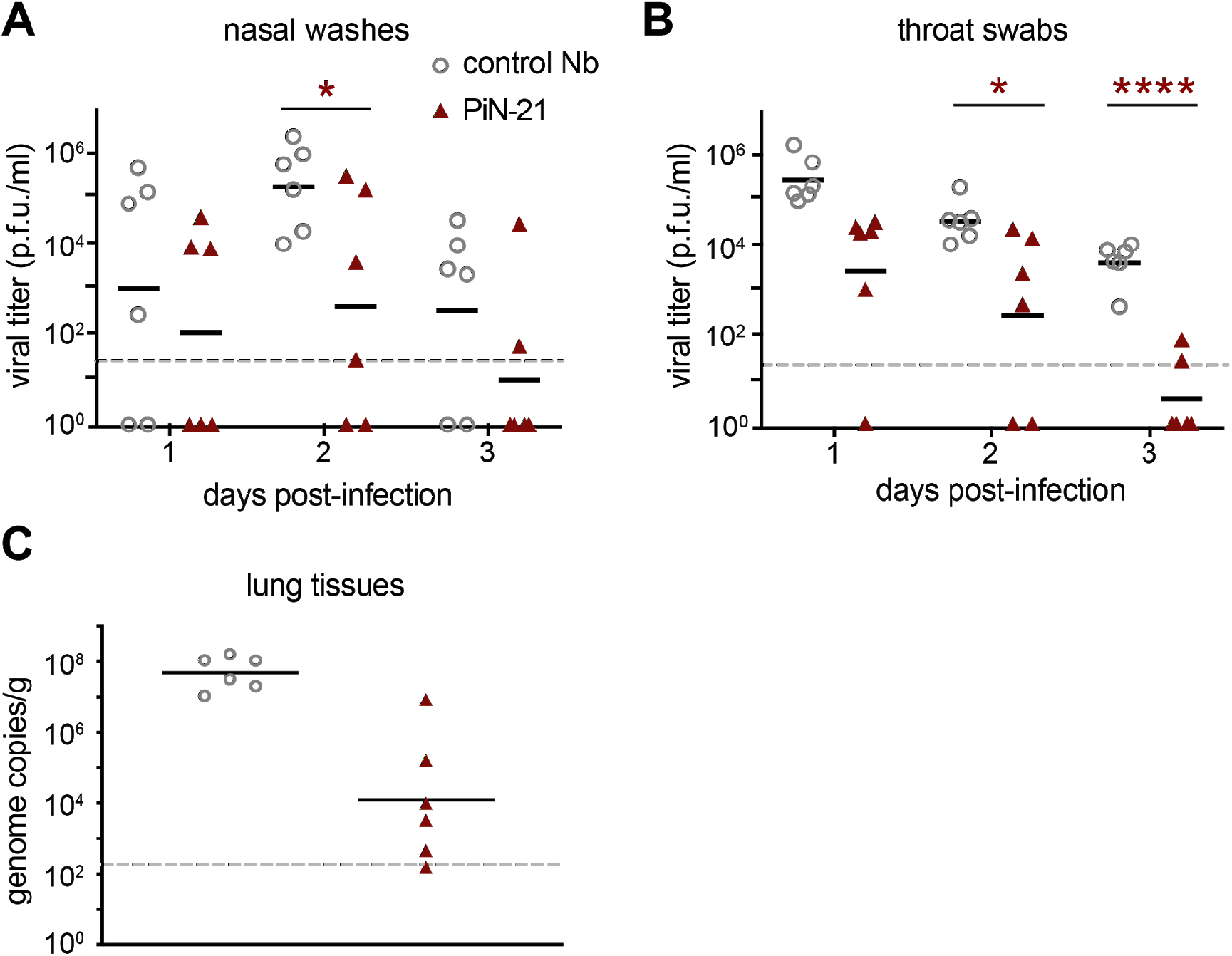
Treatment efficacy of aerosolized PiN-21 in the Syrian hamster model of SARS-CoV-2 infection. **A-B.** Measurement of viral titers in nasal washes and throat swabs using the plaque assay. Significant differences were denoted using *, P < 0.05, ****, P < 0.0001. **C.** Measurement of gRNA in lungs by RT-qPCR on 3 d.p.i. (n = 6).

**Figure S5.**
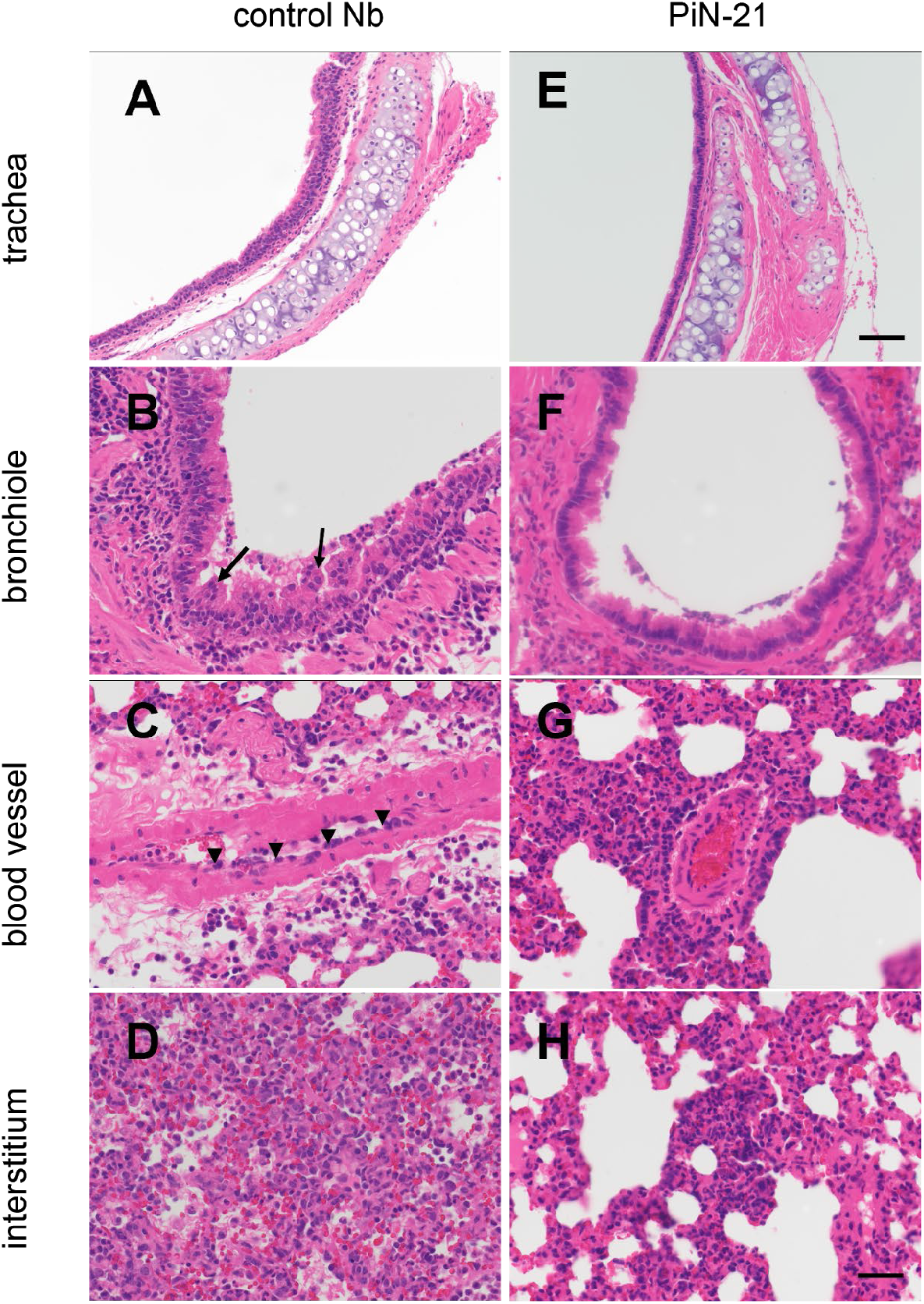
Aerosolization treatment of PiN-21 in SARS-CoV-2 infected Syrian hamsters prevents severe histopathological manifestations in the LRT. Hematoxylin and Eosin stain. Total magnification 200x, scale bar = 100 μm (**A & E**) or 400x, scale bar = 50 μm (**B,C,D,F,G,H**) **A.** Tracheal hyperplasia and hypertrophy. **B.** Bronchiole hyperplasia, degeneration, and necrosis with syncytial cells (arrows). **C.** Perivascular edema and inflammation infiltrate with reactive endothelium (arrowheads). **D.** Severe interstitial pneumonia with intra-alveolar fibrin and hemorrhage. **E.** Histologically normal trachea. **F.** Mild bronchiole degeneration with denuded intraluminal epithelium. **G.** Mild perivascular mononuclear infiltrate. **H.** Mild focal interstitial pneumonia.

**Figure S6.**
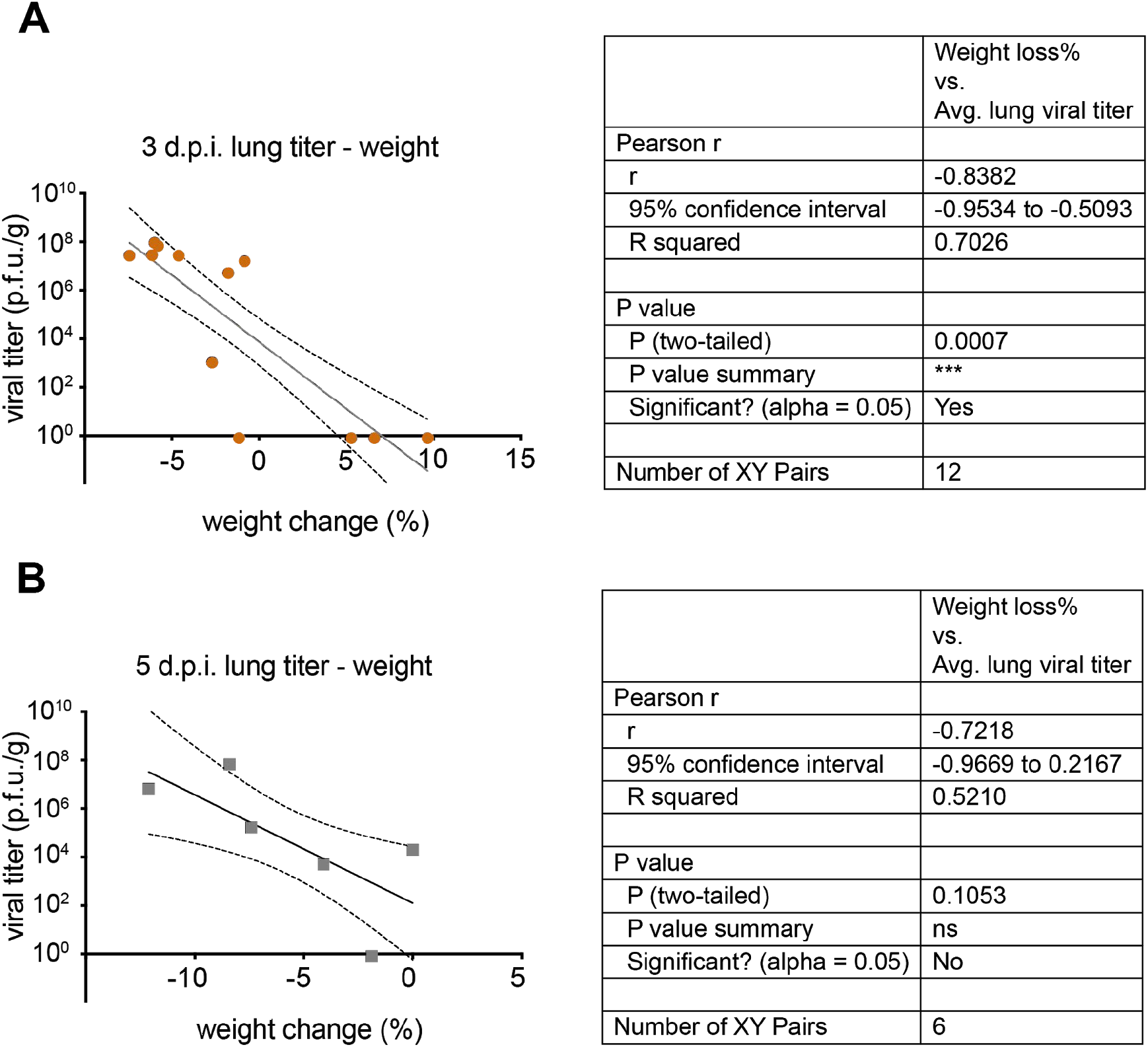
Correlation analysis of weight loss with virus titer in the hamster model. **A.** Correlation analysis on 3 d.p.i. **B.** Correlation analysis on 5 d.p.i.

## Methods and Materials

### Ethics

The animal work performed adhered to the highest level of humane animal care standards. The University of Pittsburgh is fully accredited by the Association for Assessment and Accreditation of Laboratory Animal Care (AAALAC). All animal work was performed under the standards of the Guide for the Care and Use of Laboratory Animals published by the National Institutes of Health (NIH) and according to the Animal Welfare Act guidelines. All animal studies adhered to the principles stated in the Public Health Services Policy on Humane Care and Use of Laboratory Animals. The University of Pittsburgh Institutional Animal Care and Use Committee (IACUC) approved and oversaw the animal protocols for these studies (#20067405).

### Biological safety

All work with SARS-CoV-2 was conducted under biosafety level-3 (BSL-3) conditions in the University of Pittsburgh Center for Vaccine Research (CVR) and the Regional Biocontainment Laboratory (RBL). Respiratory protection for all personnel when handling infectious samples or working with animals was provided by powered air-purifying respirators (PAPRs; Versaflo TR-300; 3M, St. Paul, MN). Liquid and surface disinfection was performed using Peroxigard disinfectant (1:16 dilution), while solid wastes, caging, and animal wastes were steam sterilized in an autoclave.

### Nanobody production

The PiN-21 construct was synthesized from Synbio Biotechnologies. Nb21_Alb_ and PiN-21_Alb_ were generated by sub-cloning a human serum albumin binding Nb (*39*) into the N-terminus of Nb21 and PiN-21 constructs (*28*). The plasmid was transformed into BL21 (DE3) cells and plated on LB-agar with 50 μg/ml ampicillin at 37°C overnight. Single bacterial colonies were picked and cultured in LB broth to reach an O.D. of ~ 0.5-0.6 before IPTG induction (0.5 mM) at 16°C overnight. Cells were then harvested, sonicated, and lysed on ice with a lysis buffer (1xPBS, 150 mM NaCl, 0.2% TX-100 with protease inhibitor). After cell lysis, his-tagged Nbs were purified by Cobalt resin and natively eluted using the imidazole buffer. Eluted Nbs were subsequently dialyzed into 1x DPBS, pH 7.4. For animal studies, endotoxin was removed with the ToxinEraser™ Endotoxin Removal Kit (Genscript), and the endotoxin level was measured using the ToxinSensor™ Chromogenic LAL Endotoxin Assay Kit (Genscript) to make sure <1 EU/ml. The proteins were sterile-filtered using the 0.22 μm centrifuge filters (Costar) before use.

### Virology

The SARS-CoV-2 isolate used was a passage 3 (p3) of the Munich isolate described previously (*48*). The virus was titrated by plaque assay and titers are expressed as plaque-forming units (p.f.u.) per ml.

### General animal procedures

Syrian hamsters (aged 3-6 months old both male and female) were obtained from the Charles River, MA. For procedures (virus infection, throat swab, and nasal wash collection), each animal was sedated with 3-5% Isoflurane. Baseline body weights were measured for all animals before infection. The animals were monitored twice daily for signs of COVID-19 disease (ruffled fur, hunched posture, labored breathing, anorexia, lethargy) post-challenge with SARS-CoV-2. Bodyweight was measured once daily during the study period. At necropsy, small pieces of the lung were collected for viral load determination. Throat swabs were collected using ultrathin swabs (Puritan™ PurFlock™ Ultra Sterile Flocked Swabs) which were placed in Opti-MEM (Invitrogen) containing double strength Antibiotic-Antimycotic (anti-anti; Life technologies). Nasal washes were collected using 500 μl of PBS with anti-anti. All samples were stored at −80°C until viral load determination. The whole trachea and lungs were collected in Opti-MEM, triazole, or 4% PFA respectively for virus titrations, RT-qPCR, and histopathological examinations.

Exp 1019: Under Isoflurane anesthesia, hamsters were infected (300 μl) with 9 x 10^4^ p.f.u. (300 μl) of SARS-CoV-2 via IT administration, immediately followed by IN administration of 100 μg (50 μl per nare) PiN-21 or a control Nb.

Exp 2021: Under Isoflurane anesthesia, hamsters were infected (50 μl per nare) with 3 x 10^4^ p.f.u. of SARS-CoV-2. At 6 h.p.i., animals were administered with 100 μg (50 μl per nare) PiN-21 (n=6) or a control Nb (n=6).

Exp 3: Hamsters were administered with PiN-21 and PiN-21_Alb_ via aerosol route and sacrificed at 8 (n=3) and 24 h (n=3) post-administration.

Exp 4: Under Isoflurane anesthesia, hamsters were infected intranasally (50ul per nare) with 3 x 10^4^ p.f.u. of SARS-CoV-2. At 6 h.p.i., animals were administered with PiN-21 (n=6) or a control Nb (n=6) *via* the aerosol route.

### Bronchoalveolar lavage (BAL) collection

Lungs with trachea were harvested from euthanized animals. A Sovereign Feeding Tube (Covetrus) was cut to the optimal length and connected to a 5 ml syringe (BD) containing 3 ml PBS with anti-anti before placement into the trachea. The PBS was gently pushed into the lungs until they were fully inflated after which the liquid (BAL) was drawn back into the syringe.

### Sample extraction and processing

For tissues, 100-200 mg of tissue was harvested, suspended in 1 ml Opti-MEM supplemented with 2X anti-anti, and homogenized using a D2400 homogenizer (Benchmark Scientific). The eluate from swabs and nasal washes were analyzed directly. Virus isolations were performed by inoculation of tissue homogenates (100 μl) onto Vero E6 cells (Hartman et al, 2020). For the preparation of RNA, tissue homogenate, swab eluate, or nasal wash (100 μl) was added to 400 μl of Trizol LS (Ambion) and thoroughly mixed by vortexing. To ensure virus inactivation, the samples were incubated for 10 minutes at room temperature and stored overnight at −80°C prior to removal from the BSL-3 facility. Subsequent storage at −80°C or RNA isolation and one-step RT-qPCR analyses were performed at BSL-2. RNA was extracted from these samples using Direct-zol RNA purification kits (Zymo Research) according to the manufacturer’s instructions. Viral RNA was detected by RT-qPCR targeting the SARS-CoV-2 nucleocapsid (N) segment as previously described (*48*). Data were normalized by tissue weight and are reported as copies of RNA determined by comparing the cycle threshold (C_T_) values from the unknown samples to C_T_ values from a positive-sense SARS-CoV-2 vRNA standard curve as previously described (*48*). Graphs were generated using GraphPad Prism, version 9.

### Neutralization assay

Nbs or hamster serum dilution (100 μl) was mixed with 100 μl of SARS-CoV-2 (Munich: P3 virus) containing 75 p.f.u. of the virus in Opti-MEM. The serum–virus mixes (200 μl total) were incubated at 37°C for 1 h, after which they were added dropwise onto confluent Vero E6 cell monolayers in six-well plates. After incubation at 37°C, 5% (v/v) CO_2_ for 1 h, 2 ml of 0.1% (w/v) immunodiffusion agarose in DMEM supplemented with 10% (v/v) FBS and 2× anti-anti was added to each well. After incubation at 37°C, 5% (v/v) CO_2_ for 72 h, the agarose overlay was removed and the cell monolayer was fixed with 1 ml/well formaldehyde [37% (w/v) formaldehyde stabilized with 10–15% (v/v) methanol] for 20 min at room temperature. Fixative was discarded and 1 ml/well 1% (w/v) crystal violet in 10% (v/v) methanol was added. Plates were incubated at room temperature for 20 min and rinsed thoroughly with water. Plaques were then enumerated and the 80% and/or 50% plaque reduction neutralization titer (PRNT_80_ or PRNT_50_) was calculated (*28, 48*). Controls of a validated SARS-CoV-2 antibody-negative, positive human serum, and an uninfected cell, were performed to ensure that virus neutralization was specific.

### Nanobody aerosolization

Aerosol exposures of hamsters to nanobodies were performed under the control of the Aero3G aerosol management platform (Biaera Technologies, Hagerstown, MD) as previously described for rodents (*49*). Hamsters were loaded into metal exposure cages and transported via mobile transfer cart to the Aerobiology suite in the RBL. There they were transferred into a class III biological safety cabinet and placed inside a rodent whole-body exposure chamber. Hamsters were exposed for 12-15 minutes to small particle aerosols containing nanobodies generated by the Aerogen Solo vibrating mesh nebulizer (Aerogen, Chicago, IL)(*19*). The system was set in a push/pull configuration with an equal volume of input air (19.5 liters per minute (lpm) total: 7.5 lpm generator, 12 lpm dilution air) and exhaust (19.5 lpm total: 6 lpm sampler, 5 lpm particle sizer, 8.5 additional vacuum) equal to 0.5 air changes/minute in the exposure chamber. To determine inhaled dose, an all-glass impinger (AGI; Catalog # #7541-10, Ace Glass, Vineland, NJ) containing 10 ml of PBS + 0.001% antifoam was attached to the chamber and operated at 6 lpm, −6 to −15 psi. Particle size was measured once during each exposure at 5 minutes using an Aerodynamic Particle Sizer (TSI, Shoreview, MN) operating at 5 lpm. A 5-minute air wash followed each aerosol, after which animals were returned to their cage. AGI samples were evaluated to determine the concentration of nanobodies recovered from the aerosol. The inhaled dose was determined as the product of the nanobody aerosol concentration, duration of exposure, and the minute volume of the individual hamster (*50*). Minute volume was determined using Guyton’s formula (*51*).

### Histologic processing and analysis

Tissue samples were fixed for a minimum of 24 h in 4% PFA before being removed from BSL-3 and subsequently processed in a Tissue-Tek VIP-6 automated vacuum infiltration processor (Sakura Finetek) and embedded in paraffin using a HistoCore Arcadia paraffin embedding machine (Leica). 5 μm tissue sections were generated using an RM2255 rotary microtome (Leica) and transferred to positively charged slides, deparaffinized in xylene, and dehydrated in graded ethanol. Tissue sections were stained with hematoxylin and eosin for histologic examination, with additional serial sections utilized for immunohistochemistry (IHC). A Ventana Discovery Ultra (Roche) tissue autostainer was used for IHC. Specific protocol details are outlined in **Table S5-6**. The histomorphological analysis was performed by a single board-certified veterinary pathologist (N.A.C.), who developed an ordinal grading score encompassing the diversity and severity of histologic findings using isotype control administered animals as a baseline. Histologic criteria were broken down into three compartments: airways, blood vessels, and interstitium, with results utilized to generate a cumulative lung injury score. This score also incorporated the overall degree of immunoreactivity to the SARS-CoV-2 S antigen. A summary of individual animal scores and specific criteria utilized to score lungs is included in **Tables S2-3**.

### Multispectral Whole Imaging

Brightfield and fluorescent images were acquired using a Mantra 2.0™ Quantitative Pathology Imaging System (Akoya Biosciences). To maximize signal-to-noise ratios, fluorescent images were spectrally unmixed using a synthetic library specific for the Opal fluorophores used for each assay and for DAPI. An unstained Syrian hamster lung section was used to create an autofluorescence signature that was subsequently removed from images using InForm software version 2.4.8 (Akoya Biosciences).

### Nanobody aerosolization using the mesh nebulizer

Nb (Nb21_Alb_, PiN-21 and PiN-21_Alb_) was concentrated to 1 ml (1.5 mg/ml) in 1xDPBS. 0.5 ml was saved as a control for ELISA and pseudovirus neutralization assay. The other 0.5 ml was aerosolized by using a portable mesh atomizer nebulizer (MayLuck). No obvious dead volume was observed. The aerosolized droplets were collected in a microcentrifuge tube. The concentration was measured to calculate the recovery of the proteins.

### Pseudotyped SARS-CoV-2 neutralization assay

Pseudotype neutralization assay was carried out and IC50 was calculated as previously described (*28*).

## Contributions

Y.S. and W.P.D. conceived and supervised the study. W.B.K. oversaw the study. S.N. performed the animal studies and virology experiments. Y.X. and N.L.T. helped S.N. in animal studies. Y.X. performed Nb purification, molecular biology and biochemistry. Y.X. and N.L.T. performed viral RNA analysis. L.J.R. performed tissue culture and helped in viral RNA analysis. D.S.R. carried out the aerosolization of Nbs. N.A.C. designed and optimized immunohistochemical assays and carried out the histopathologic analysis. Y.S., W.P.D., S.N., and Y.X. design the experiments and interpret the results. Y.X., Y.S., S.N., and N.A.C. prepared the figures. Y.S. drafted the manuscript with edits from Y.X., S.N., N.A.C., and W.P.D.

## Acknowledgments

We would like to thank Hans P. Gertje (histology), Katherine O’Malley (aerosol), Morgan Midgett (aerosol), Matthew D. Dunn (animal studies), Theron Gilliland (animal studies), Emily L. Cottle (animal studies), Reagan C. Walker (animal studies), Stacey R. Barrick (animal studies) and Kuntal A. Thakkar (laboratory) for excellent technical assistance.

